# Global Modulation in DNA Epigenetics during Pro-Inflammatory Macrophage Activation

**DOI:** 10.1101/587311

**Authors:** Nikhil Jain, Tamar Shahal, Tslil Gabrieli, Noa Gilat, Dmitry Torchinsky, Yael Michaeli, Viola Vogel, Yuval Ebenstein

## Abstract

DNA methylation patterns create distinct gene expression profiles. These patterns are maintained after cell division, thus enabling the differentiation and maintenance of multiple cell types from the same genome sequence. The advantage of this mechanism for transcriptional control is that chemical-encoding allows to rapidly establish new epigenetic patterns “*on-demand*” through enzymatic methylation and de-methylation of DNA. Here we show that this feature is associated with the fast response of macrophages during their pro-inflammatory activation. By using a combination of mass spectroscopy and single-molecule imaging to quantify global epigenetic changes in the genomes of primary macrophages, we followed three distinct DNA marks (methylated, hydroxymethylated and unmethylated), involved in establishing new DNA methylation patterns during pro-inflammatory activation. The observed epigenetic modulation together with gene expression data generated for the involved enzymatic machinery, may suggest that de-methylation upon LPS-activation starts with oxidation of methylated CpGs, followed by excision-repair of these oxidized bases and their replacement with unmodified cytosine.

## Introduction

Macrophages are an integral component of our host immune response. During several pathological conditions, uncommitted macrophages (M0) get activated into a spectrum of states, from the pro-inflammatory (M1) to pro-healing (M2) phenotypes with gradually altered transcriptional profiles and functional outputs (1, 2). Many molecular mechanisms of these activation processes, including the underlying biochemical pathways and the involved transcription factors and co-factors, have been extensively characterized.

With the advent of new tools and techniques for genomic analysis, it has been postulated that epigenetic modifications are critical for establishing efficient gene-expression profiles during inflammation and disease (3, 4). Accordingly, alteration of the genomic methylation pattern may be critical to the fast response of macrophages to chemical and mechanical changes in their microenvironment. Investigations of the epigenetic changes during macrophage activation have largely been focused on modification of histones (5, 6). A spectrum of different histone modifications has been reported, which can be broadly divided into marks that promote or suppress transcription. For example, a repressed state is characterized by the presence of marks like histone 3 lysine 9 trimethylation (H3K9me3) and H3K27me3, whereas an activated state is marked with the presence of histone marks like H3K4me3 and H3K9/14-Ac (4, 6, 7).

Despite extensive characterization of histone-modification changes during macrophage activation, epigenetic changes involving DNA modifications have largely remained unexplored. DNA methylation is a central epigenetic marker modulated during development, differentiation and disease. Methylation of the 5-carbon of cytosine (5mC), is the most studied and among the most significant epigenetic modification. 5mC is found in all tissues and cell types, but at different levels (8). In mammalian DNA, methylation mostly occurs on cytosines within CpG dinucleotides and approximately 60% of human gene promoters contain clusters of CpGs referred to as CpG islands. CpG methylation is regulated by a family of enzymes called DNA methyltransferases (DNMTs) that catalyze the transfer of a methyl group from the methyl group donor S-adenosyl-L-methionine to the fifth carbon of cytosine residues (9). CpG methylation is generally associated with gene silencing by affecting the recruitment of transcription factors. Thus, the methylation status of gene promoters may predict gene activity and relate epigenetic transformations to gene expression and protein abundance.

Recently, more attention has been given to 5-hydroxymethylcytosine (5hmC) (10), which is an oxidation product of 5mC catalyzed by the ten-eleven-translocation (Tet) proteins in the process of de-methylation (11-13). 5hmC is found in all tissues and cell types investigated to date (14, 15) and similar to 5mC is almost exclusively found at CpGs (16, 17). However, contrary to 5mC, the presence of 5hmC has generally been associated with increased gene expression (18-21). Loss of DNA methylation *via* 5hmC has been observed in different biological contexts and this alteration can take place actively or passively. Active DNA demethylation is an enzymatic process that removes oxidized 5mC residues through Thymine DNA glycosylase (TDG)-mediated base excision repair. By contrast, passive DNA demethylation refers to loss of 5mC after initial oxidation to 5hmC followed by successive rounds of replication. 5hmC has no known maintenance mechanism and is eventually replaced by cytosine after replication (22).

Emerging evidence suggests a critical role of these DNA modifications in modulating transcription profiles during macrophage activation and reprogramming (23), but existing data are limited to modifications in the vicinity of specific genes (24, 25). The redistribution of 5hmC has been studied during differentiation of monocytes into macrophages (26) but global epigenetic changes for macrophage activation during immune tolerance and innate immune have largely been unexplored. It is still largely unclear whether the pro-inflammatory activation of macrophages is associated with global changes in 5mC and 5hmC levels. Understanding such epigenetic changes is necessary for characterizing the behaviour of macrophages during homeostasis and pathological conditions like chronic inflammation and disease.

Commonly used methods for studying 5mC methylation include HPLC coupled to mass spectrometry (27), immuno-based assays like dot-blot (28), immunohistochemical assays (29) and enzyme-linked immunosorbent assays (30, 31). However, most of these techniques suffer from shortcomings such as low sensitivity and poor detection limits, high costs and the requirement of large amounts of starting material. The low levels of 5hmC in macrophages make these cells exceptionally challenging for quantitative analysis. Hence, in this work we used two recently developed methods which utilizes an ultra-sensitive single-molecule approach by which fluorescently-labeled 5mC (32) and 5hmC (33) marks can be directly visualized and counted on individual chromosomal fragments(34). Using these methods, we find that the pro-inflammatory (M1) activation of macrophages with lipopolysaccharide (LPS) leads to a global reduction in 5mC levels accompanied by an increase in both 5hmC levels and the amount of unmethylated CpGs. These results suggest a significant reduction in total DNA methylation within 24 h of LPS stimulation. Additionally, this process involves conversion of 5mC to 5hmC and restoration of unmodified CpGs during macrophage M1 activation. Since 5mC levels largely depends on the activity of DNA Methyl Transferases (DNMTs) and levels of 5hmC depend on the activity of Ten-eleven Translocation (Tet) enzymes, we also measured the expression levels of various DNMTs and Tet enzymes, which were found to be downregulated and upregulated upon LPS stimulation, respectively. Furthermore, increased expression levels of the Thymine-DNA Glycosylase (TDG) enzyme, together with the increase in unmethylated CpGs suggest active loss of DNA methylation during macrophage activation, which is mediated by oxidation and excision-repair of methylated cytosines.

## Methods

### Bone Marrow Isolation, Macrophage Differentiation and Culture

Femurs were isolated from healthy mice and bone marrow was flushed with PBS and passed through 7 μm cell strainer to obtain single cell suspension. Bone marrow derived macrophages (BMDM) were centrifuged, suspended in BMDM culture media and 2 million macrophages were seeded in 60 mm non-treated plastic dishes for 7 days. Equal volume of media was again added on day 4. Cells were used on day 7 for further experiments. BMDM culture media contains high glucose DMEM, 10%FBS, Glutamine, Non-essential amino acids, Sodium pyruvate, Beta-mercaptoethanol, Penicillin-Streptomycin and L929 media. L929 media was prepared by culturing 2 million L929 cells in 200 ml media in a 300 cm^2^ flask for 8 days without changing media. The media used for L929 cell culture contains RPMI, 10% FBS, Glutamine, Non-essential amino acids, Sodium pyruvate, HEPES and Beta-mercaptoethanol. For M1 activation, macrophages were activated with LPS (25 ng/ml) for 24 h.

### DNA Isolation

DNA from Control (M0) and 24 h LPS activated (M1) macrophages was purified in agarose plugs in order to maintain large DNA fragments, following the Bionano Genomics IrysPrep protocol with slight modifications. Briefly, 1 × 10^6^ cells were washed twice with PBS, resuspended in cell suspension buffer (CHEF mammalian DNA extraction kit, Bio-Rad), and incubated at 43 °C for 10 min. 2% low melting agarose (CleanCut agarose, Bio-Rad) was melted at 70 °C followed by incubation at 43 °C for 10 min. Melted agarose was added to the resuspended cells at a final concentration of 0.7% and mixed gently. The mixture was immediately cast into a plug mold, and plugs were incubated at 4 °C until solidified. Plugs were incubated twice (2 h of incubation followed by an overnight incubation) at 50 °C with 167 μL of freshly prepared proteinase K (Qiagen) in 2.5 mL of lysis buffer (BioNano Genomics Inc.) with occasional shaking. Next, plugs were incubated with 50 μL of RNase (Qiagen) in 2.5 mL of TE (10 mM Tris, pH 8, 1 mM EDTA) for 1 h at 37 °C with occasional shaking. Plugs were washed three times by adding 10 mL of wash buffer (10 mM Tris, pH 8, 50 mM EDTA), manually shaking for 10 s, and discarding the wash buffer before adding the next wash. Plugs were then washed four times by adding 10 mL of wash buffer and shaking for 15 min on a horizontal platform mixer at 180 rpm at room temperature. Plugs were then washed three times in TE (pH 8) and were melted for 2 min at 70 °C, followed by 5 min of incubation at 43 °C. Next, 0.4 units of Gelase (Epicenter) were added and the mixture was incubated for 45 min. High-molecular-weight DNA was purified by drop dialysis using a 0.1 μm dialysis membrane (Millipore) floated on TE (pH 8). Viscous DNA was gently pipetted and incubated at room temperature overnight in order to achieve homogeneity. DNA concentration was determined using Qubit BR dsDNS assay (Thermo Fisher Scientific).

### 5mC quantification by mass spectroscopy (Assay 1)

The analysis of the percent of 5mC per total nucleotides, by LC-MS/MS was conducted according to the detailed procedure described before (35). Briefly, macrophage DNA samples were hydrolyzed to nucleosides by a combination of three enzymes: Nuclease S1, antarthicphosphetase and phosphodiesterase. The chromatographic separations were performed on an Acquity UPLC system (Waters) using Xselect HSS T3 column. Nucleosides were eluted with a gradient of 0.1% formic acid in water and 0.1% formic acid in acetonitrile and directed into a Xevo TQD triple quadrupole mass spectrometer (Waters). The measurements of targeted nucleosides were conducted in a positive ion mode by multiple reaction monitoring (MRM). The relative content of 5mC in each sample was calculated from the MRM peak area of 2’-deoxy-5’-methylcytidine (5mdC) divided by the sum of peak areas of all other nucleosides (Fig. 1A).

**Figure 1.**
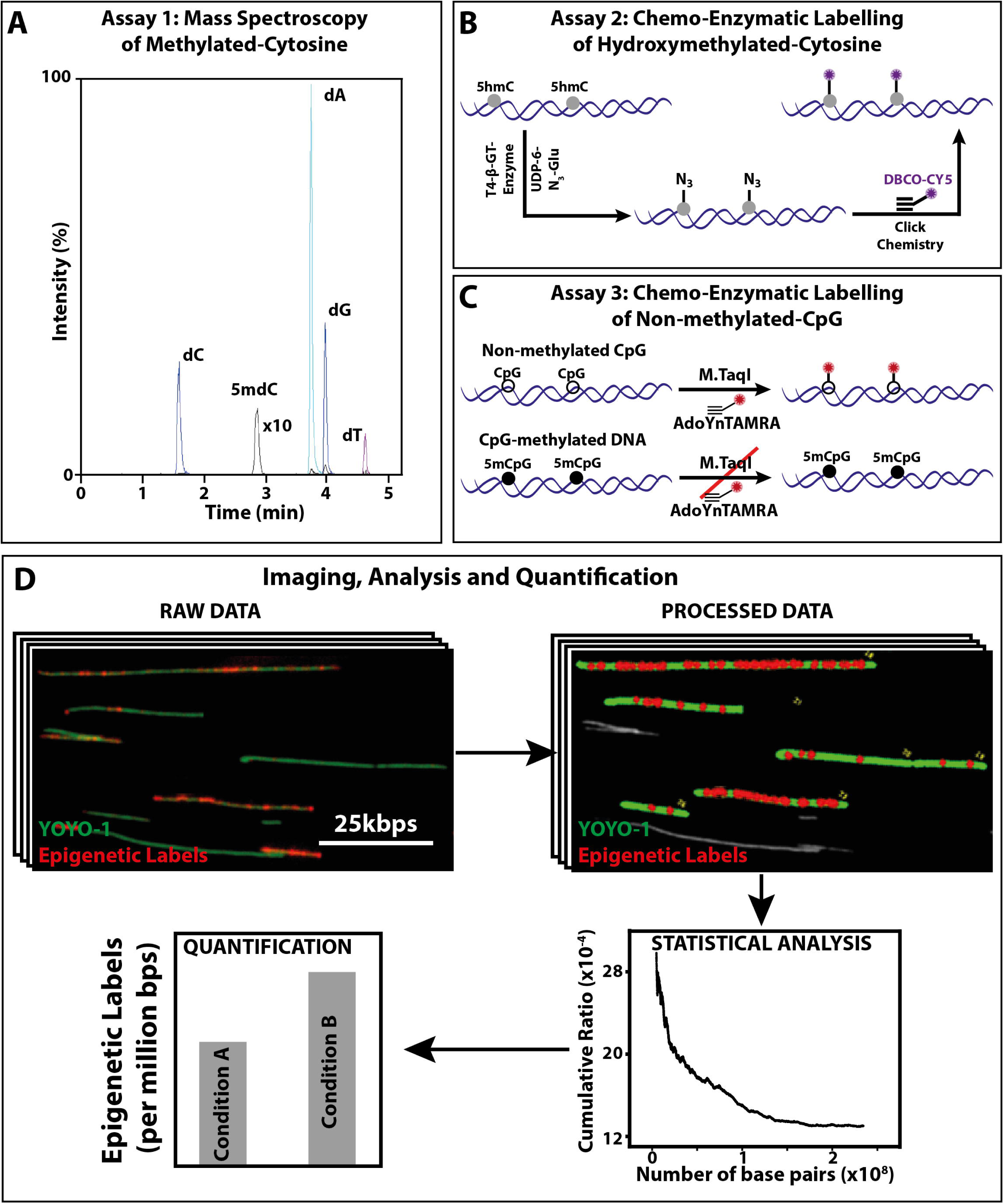
Novel optical imaging techniques to quantify 5hmC and 5mC levels of DNA: **(A) Assay-1:** An MRM chromatogram of nucleosides obtained by hydrolysis of gDNA. **(B) Assay 2:** Schematic representation of the two-step 5hmC labelling reaction. First, T4 β-GT enzymatic glucosylation of 5hmC with UDP-6-N3-Glu is performed. Next, click reaction between the N3 group and the fluorescently labelled alkyne DBCO-Cy5 is performed. This two-step reaction results in fluorescently labelled 5hmC. **(C) Assay 3:** Schematic representation of M.TaqI mediated labelling of non-methylated cytosines: Top: M.TaqI catalyzes the transfer of a TAMRA fluorophore from the cofactor AdoYnTAMRA onto the adenine residue that lies in its TCGA recognition site. Bottom: If the cytosine residue that lies within M.TaqI’s recognition site is methylated, the reaction is blocked. **(D)** Multiple fluorescence microscope images are obtained per sample, showing both YOYO-1 labelling of the entire DNA molecule (green) and the epigenetic-labels (red dots). In-house developed user interface of the image processing software, where the length of the DNA molecules is measured (green) and colocalized epigenetic-labels (red dots) are detected and subsequently quantified relative to DNA length. Output of the statistics tool, showing the total length of DNA (in bps) and the percentage of epigenetic-labels relative to DNA. Dummy comparison of the number of epigenetic labels between two conditions.

### 5hmC labeling procedure (Assay 2)

Quantification of 5hmC was performed according to our recently published protocols (19, 33, 36). Briefly, 5hmC was fluorescently labeled in a two-step chemical labeling procedure (Fig. 1B). Three hundred nanograms of genomic DNA was mixed with 3 μl of buffer 4 (NEB), 0.5 μl of UDP-6-N3-Glu (0.3 mM), 2 μl of T4 bacteriophage β-glucosyltransferase (T4-BGT, NEB) and ultrapure water to a final volume of 30 μl. The reaction was mixed and incubated overnight at 37 °C. On the following day, 0.15 μl of DBCO-Sulfo-Cy5 (10 mM, Jena Biosciences) was added to the mixture, followed by a second overnight incubation at 37 °C (click reaction) (Fig. 1B). Upon completion of the labeling procedure, samples were washed by ethanol precipitation. Samples were stored at 4 °C until analyzed.

### Unmodified CpG labelling procedure (Assay 3)

Quantification of unmodified CpGs was performed according to our recently published protocol (32, 37). Briefly, to generate methylation sensitive labels we used the DNA MTase M.TaqI, which catalyzes the transfer of a carboxytetramethylrhodamine (TAMRA) fluorophore from the synthetic cofactor AdoYnTAMRA onto the adenine residue within its recognition sequence (TCGA). M.TaqI enzyme can distinguish methylated from non-methylated cytosines (Fig. 1C), and has been shown recently that it can be used to detect and quantify methylation levels in DNA molecules. The bacterial methyltransferase M.TaqI in combination with a synthetic cofactor analogue can fluorescently label the adenine residue in TCGA sites containing non-methylated CpGs. M.TaqI has potential access to only about 5.5% of all CpGs, however, these sites are present in over 90% of gene promoters.

The labelling reaction was carried out as follows: 300 ng of DNA isolated from Control and LPS-stimulated macrophage was reacted with 0.5 μl of M.TaqI, 2 μl of Cut-smart buffer and 40 nM of AdoYnTAMRA in labelling buffer (20 mM Tris-HOAc, 10 mM Mg(Cl)2, 50 mM KOAc, 1 mM DTT, pH 7.9) in a total reaction volume of 20 μl at 60 °C for 1 h. The labelled DNA was reacted with 3 μl (40 μg) of proteinase K (Sigma) at 45°C for 1 hour to disassemble protein-DNA aggregates. Post labelling, DNA was cleaned by ethanol precipitation. Prior to imaging, the labelled DNA was stained with 0.5 μM of YOYO-1 (Invitrogen) for visualization of its contour. DTT (Sigma) was added to the reaction (200 mM) to prevent photo bleaching and DNA breaks (32) (Fig. 1C).

### Preparation of activated glasses for stretching

Glass surfaces for DNA extension were prepared as (38) with minor modifications. In short, standard glass cover slides were first cleaned by immersing them overnight in 2:1 (v/v) mixture of nitric acid (70%) and hydrochloric acid (37%), respectively. On the following day, the slides were washed extensively by using ultrapure water and ethanol and dried under nitrogen stream. Dried slides were inserted into a 300 ml aqueous solution containing 1.2 ml N-trimethoxysilylpropyl-N,N,Ntrimethylammonium chloride (Gelest) and 180 μl vinyltrimethoxysilane (Gelest), then incubated for 17.5 h at 65 °C with mild shaking. Slides were then washed extensively by using ultrapure water and ethanol and stored in ethanol at 4 °C until use (33).

### Single molecule imaging

Labeled samples were diluted 1:100–1:300 in HEPES DTT buffer (100 mM HEPES, 200 mM DTT, Sigma) and were stained with 0.5 μM of YOYO-1 (Invitrogen) DNA intercalating dye. 8 μl of each sample were extended on a glass slide by placing the solution at the interface of an activated coverslip and a clean microscope slide. Labeled samples were then stretched on glass slides and imaged using a fluorescence microscope. The extended DNA molecules were imaged with a fluorescence microscope (TILL photonics GmbH) using an Olympus UPlanApo 100X 1.3 NA oil immersion objective. Images were captured with the appropriate filters (485/20ex and 525/30em, 560/25ex and 607/36em, and 650/13ex and 684/24em (Semrock Inc., Rochester NY, USA) for the YOYO-1, TAMRA, and Cy5 channels respectively). Images were acquired by a DU888 EMCCD (Andor technologies) (33).

### Image processing and Data analysis

DNA molecules appear as extended green lines (Fig. 1D) dotted with red labels indicating epigenetic labels (Fig. 1D). Some DNA molecules were unlabeled, few were sparsely labeled and some were heavily labeled (Fig. 1D). Finally, images were processed, analyzed and quantified using in-house image processing software (Fig. 1D). Statistical analysis of the data was performed to plot cumulative frequency of epigenetic labels as a function of the total amount of DNA acquired (calculated as number of epigenetic labels/total DNA length in base pairs). This analysis helps in assessing the required amount of data needed for accurate methylation measurement; upon sampling more data, the labelled epigenetic site levels readout stabilizes, and finally reaching a plateau, indicating sufficient data was sampled (Fig. 1D). The total DNA length of each molecule was calculated, and the number and intensity of colocalized labels on each molecule was counted (33).

### RNA Isolation and Real-Time qPCR Analysis

RNA was isolated with Qiagen RNA Isolation Kit as per the manufacturer’s protocol. cDNA was prepared using iScript Advanced cDNA Synthesis Kit (172-5038 BioRad). Real-time qPCR experiments were performed using SsoFast EvaGreen Supermix (172-5203 Bio-Rad) in CFX96 Real-Time PCR Detection System (Bio-Rad).

### RNA sequencing

Detailed information regarding RNA sequencing is available in our recently published work (39). Briefly, RNA was isolated with Qiagen RNA Isolation Kit as per the manufacturer’s protocol. For library preparation, the quantity and quality of the isolated RNA were determined with a Qubit® (1.0) Fluorometer (Life Technologies, California, USA) and a Tapestation 4200 (Agilent, Waldbronn, Germany). The TruSeq Stranded HT mRNA Sample Prep Kit (Illumina, Inc, California, USA) was used in the subsequent steps following the manufacturer’s protocol. Briefly, total RNA samples (1 μg) were ribosome depleted and then reverse-transcribed into double-stranded cDNA with Actinomycin added during the first-strand synthesis. The cDNA samples were fragmented, end-repaired and polyadenylated before ligation of TruSeq adapters. The adapters contain the index for dual multiplexing. Fragments containing TruSeq adapters on both ends were selectively enriched with PCR. The quality and quantity of the enriched libraries were validated using Qubit® (1.0) Fluorometer and the Tapestation 4200 (Agilent, Waldbronn, Germany). The product is a smear with an average fragment size of approximately 360 bps. The libraries were normalized to 10 nM in Tris-Cl 10 mM, pH 8.5 with 0.1% Tween 20. For cluster generation and sequencing, the Hiseq 4000 SR Cluster Kit (Illumina, Inc, California, USA) was used using 8 pM of pooled normalized libraries on the cBOT V2. Sequencing was performed on the Illumina HiSeq with single end 125 bps using the HiSeq 3000/4000 SBS Kit (Illumina, Inc, California, USA). Reads quality was checked with FastQC (https://www.bioinformatics.babraham.ac.uk/projects/fastqc/) and sequencing adapters were trimmed using Trimmomatic (40). Reads at least 20 bases long, and an overall average Phred quality score greater than 10 were aligned to the reference genome and transcriptome (FASTA and GTF files, respectively, downloaded from Ensemble, genome build GRCm38) with STAR v2.5.1 with default settings for single-end reads (41). Distribution of the reads across genomic features was quantified using the R package Genomic Ranges from Bioconductor Version 3.0 (42). Differentially expressed genes were identified using the R package edgeR from Bioconductor Version 3.0 (43).

### Data Availability

RNA–seq data have been deposited in GEO. The accession codes are GSM3373977–GSM3373979 and GSM3373980–GSM3373982 for Control untreated BMDMs and LPS-treated BMDMs respectively. Molecule count data corresponding to hydroxymethylated and unmethylated CpGs on individual chromosomal DNA fragments in macrophages extracted from the images are provided as supporting information file 1&2 respectively.

### Software Availability

Image analysis software installation files, detailed step-by-step installation instructions, sample images and the corresponding image analysis file (raw) are available from GitHub (https://github.com/ebensteinLab/Tiff16_Analyzer) and labelled as Software Install, Instruction files, Examples images, and result.txt respectively.

## Results

### Pro-inflammatory activation of macrophages with LPS is marked with global changes in DNA Methylation levels

To check whether macrophage M1 activation is marked with changes in DNA methylation, bone marrow derived primary macrophages were activated with LPS for 24 h. M1 activation was confirmed by changes in cell morphology and the expression of typical gene markers (39) (Figs. 2A&B). Levels of 5mC in the two states were quantified by mass spectroscopy. We found that M1 activation of macrophages resulted in a significant decrease in total levels of 5mC (Fig. 2C). We next evaluated the levels of 5hmC in order to assess whether the reduction in 5mC occurs through Tet mediated demethylation, a process initiated by the oxidation of 5mC to 5hmC. The formation of 5hmC from 5mC inherently lowers the levels of 5mC at any given nucleotide position resulting in genome wide hypomethylation (44, 45). Fig. 2D shows that macrophage M1 activation is marked with global increase in 5hmC levels, whereby the total number of 5hmC labels were higher in DNA isolated from M1 macrophages (7.3 labels per million base pairs & supporting information file 1) as compared to M0 macrophages (4.3 labels per million base pairs & supporting information file 1).

**Figure 2.**
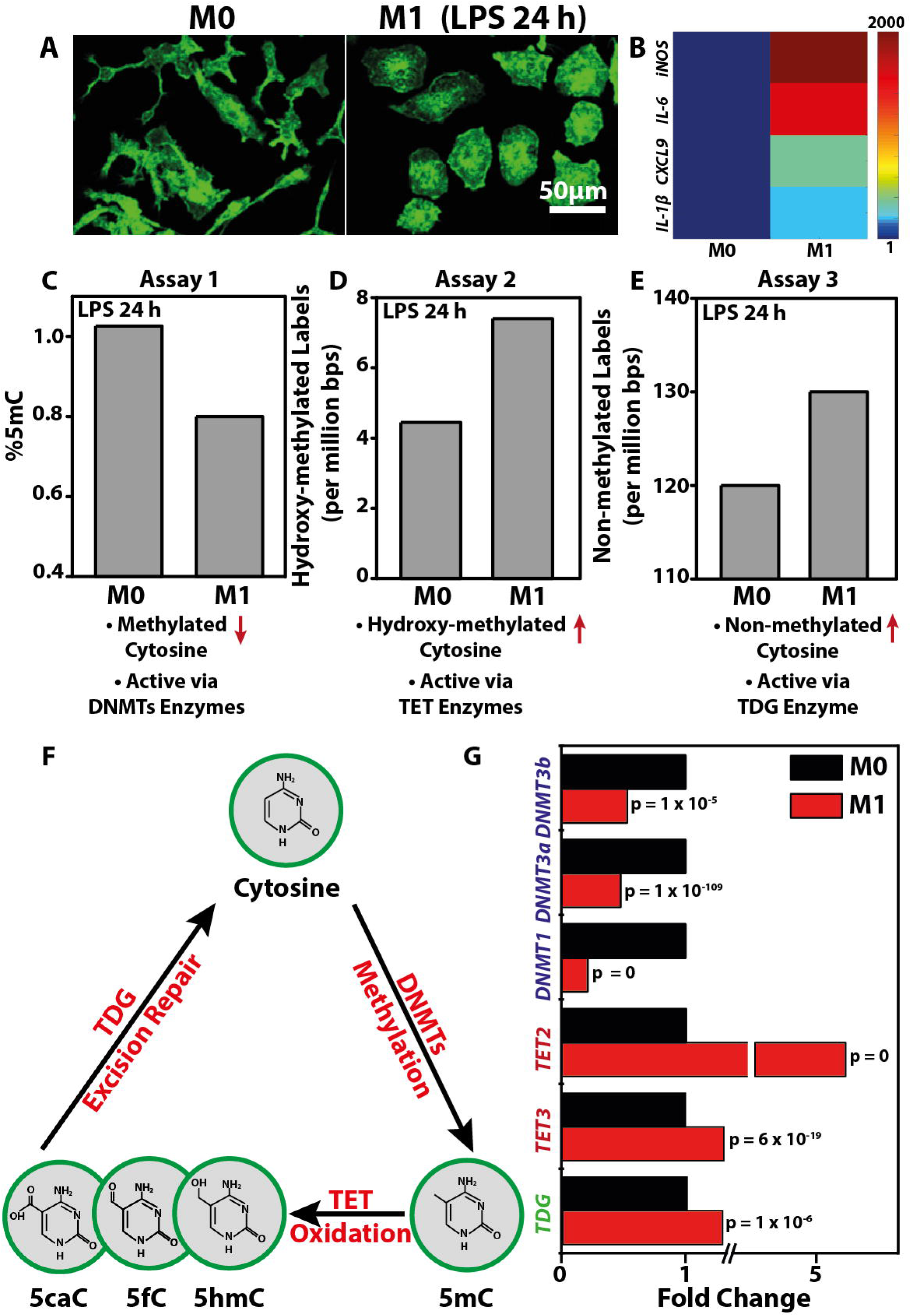
Macrophage M1 activation is marked with global changes in DNA methylation levels which is correlate with differential expression of associated enzymes: **(A)** Representative images of Control (M0) and LPS-treated (M1) (24 h) macrophages stained for F-actin (green). Scale bar = 50 μm. **(B)** Color-coded arrays show the expression levels of pro-inflammatory genes in M0 and M1 macrophages. Levels are normalized to the M0 macrophages. **(C)** Percent of 5mC of total nucleotides in M0 and M1 macrophages as obtained using Assay 1. **(D)** Bar-graph displays number of detected 5hmC labels per million base pairs as obtained using Assay 2. **(E)** Bar-graph displays number of un-methylated cytosine labels per million base pair as obtained using Assay 3. **(F)** Depiction of cytosine’s methylation and demethylation processes. The different modified forms of cytosine (5mC, 5hmC, 5fC and 5caC) along with the corresponding enzymes responsible for each modification are shown. **(G)** Bar graph shows the differential levels of various DNMTs, Tets and TDG enzymes in M0 and M1 macrophages as obtained from RNA-Seq experiment.

This is in contrast to the observations in many cancer types where both 5mC and 5hmC are globally reduced compared to healthy tissue (46). The reduction in 5hmC observed in cancer may be due to the increased proliferation rate of cancer cells that lead to rapid dilution of genomic 5hmC *via* DNA replication (47). In the case of macrophages, which are non-dividing, 5hmC residues are expected to be more persistent. Interestingly, a global reduction in DNA methylation was not only accompanied by an increase in 5hmC levels but also a significant increase in unmethylated CpG levels. M0 macrophages were found to contain less unmethylated CpGs as compared to M1 macrophages as evident from higher M.TaqI epigenetic labels in DNA isolated from M1 macrophages (Fig. 2E & supporting information file 2). This suggests that some methylated CpGs return to their native state and undergo full conversion of 5mC to cytosine. We next assessed the levels of different enzymes involved in the dynamic transformation of 5mC to 5hmC further leading to DNA demethylation, as summarized in Fig. 2F.

### Global changes in 5mC and 5hmC levels during M1 activation are associated with differential expression of related enzymes

Since lowering of 5mC has largely been attributed to a decreased expression level and activity of DNMTs, we next looked at the levels of different DNMTs during M1 activation by reanalyzing our recently published RNA-Seq data (39). DNMT1 is the most abundant DNA methyltransferase in mammalian cells and predominantly maintains methylation patterns after replication by methylation of hemimethylated CpG di-nucleotides in the mammalian genome. DNMT3, which includes DNMT3a, DNMT3b and DNMT3l, is a family of DNA methyltransferases that could *de-novo* methylate hemimethylated and unmethylated CpG and can also interact with DNMT1 (48). In line with our observation that macrophages activated with LPS for 24 h showed global reduction in 5mC levels, we also found that this reduction is accompanied with a significant reduction in the expression levels of DNMT 1, 3a and 3b (Fig. 2G). As mentioned earlier and summarized in Fig. 2F, we also checked the levels of Tet enzymes which catalyze the successive oxidation of 5mC to 5hmC, 5-formylcytosine (5fC), and 5-carboxylcytosine (5caC) (49). These 5mC oxidation products were implicated as intermediates in the conversion of 5mC to unmodified cytosines. Levels of 5hmC has recently been positively correlated with the expression levels of the Tet enzyme family (50). As shown in Fig. 2G and in line with our observation that M1 activation is marked with global increase in 5hmC levels, we found that expression levels of Tet 2 and Tet 3 increased significantly during M1 activation. This may imply that macrophage M1 activation is marked with an increase in Tet enzyme levels which in turn results in higher oxidation of 5mC to 5hmC. Lastly, we checked the levels of TDG which excises T from G·T mispairs and is thought to initiate base excision repair of deaminated 5-methylcytosine (5mC). Recent studies show that TDG is also essential for active DNA demethylation. In the case of macrophages, which are non-dividing, hence cannot passively deplete DNA methylation levels by cell division, we postulated a possible role of TDG in the global increase in unmethylated CpG levels during M1 activation. Interestingly, we found that levels of TDG enzymes were also upregulated during M1 activation suggesting an essential role of TDG in epigenetic reorganization dynamics during this inflammatory process

## Discussion

In mammals, along with other epigenetic regulators, DNA methylation is a prevalent epigenetic modification and is almost entirely found on the fifth position of the pyrimidine ring of cytosines in the context of CpG dinucleotides. 5-Methylcytosine accounts for ∼1 % of all bases, whereby the majority (∼70 %) of CpG dinucleotides throughout mammalian genomes are methylated (51). DNA methylation can have profound effects on gene expression, is crucial for proper development, and is implicated in different disease processes(52). However, the epigenetic role of DNA methylation during macrophage activation and inflammation has not been fully characterized. It is important to define how global epigenomic changes are associate with macrophage polarization and gene transcription during homeostasis and disease in order to help uncover central regulatory mechanisms.

Here, we report a significant change in the genome-wide levels of DNA modifications during macrophage M1 activation by using mass spectroscopy, and single molecule epigenetic imaging. Specifically, we recorded global reduction in methylation levels (5mC) and a concomitant increase in the levels of hydroxy-methylcytosine (5hmC) and unmodified CpGs (Figs. 2C-E). Mechanistically, we postulate these changes to be largely associated with the altered expression and hence enzyme activity of DNMTs, Tets and TDG (Fig. 2G). Altogether our work depicts a plausible active demethylation process by which 5mC is oxidized and then excised and replaced by native cytosine. In itself, this process may induce modulation in genes associated with methylation-controlled promoters or enhancers. In addition, we found that 5hmC levels are increased after 24 hours, possibly indicating an active role of 5hmC in M1 activation (Fig. 2D). It has been shown that 5hmC levels in gene bodies are positively correlated with gene expression (53). 5hmC may be involved in transient gene expression necessary for the activation process or in permanent upregulation of M1 functional genes. Our work now indicates that global changes in DNA modifications occur (Fig. 2) and can thus serve as additional biomarker for pro-inflammatory activation. We also identified possible underlying molecular intermediates involved in this dynamic regulation of epigenetic modifications during M1 activation (Fig. 2E). A genome-wide map of epigenetic evolution throughout macrophage activation and phenotypic establishment may shed light on the specific genes and non-coding regions involved in this highly dynamic process. Characterizing and targeting such epigenetic processes, as well as identifying epigenetic profiles along with the molecular intermediates that underlie disease state, may not only help in understanding the basic biology of macrophage polarization and their functional diversity, but also offers new opportunities for personalized medicine, and innovative new approaches for the treatment of inflammatory diseases.

## Acknowledgements

We want to thank Elmar Weinhold RWTH Aachen University for providing methylation reagents. This work is supported by SNF NCCR ‘Molecular Systems Engineering and SNF CR32I3_156931 grants to V.V, SNF IZK0Z3_174260 and EMBO grants to N.J, and the BeyondSeq consortium [EC program 634890] the European Research Councils starter grant [337830] and the i-Core program of the Israel Science Foundation [1902/12] to YE.

## Authors contribution

NJ, VV and YE conceived of the project and designed the experiments. TS performed the LC-MS experiments. NJ and TG did the methylation studies. NJ and NG performed the hydroxymethylation studies. NJ, DT and YE analyzed the data. NJ, VV and YE wrote the manuscript. All the authors edited the manuscript.

